# RSC primes the quiescent genome for hypertranscription upon cell cycle re-entry

**DOI:** 10.1101/2021.01.28.428695

**Authors:** Christine E. Cucinotta, Rachel H. Dell, Keean C.A. Braceros, Toshio Tsukiyama

## Abstract

Quiescence is a reversible G_0_ state essential for differentiation, regeneration, stem cell renewal, and immune cell activation. Necessary for long-term survival, quiescent chromatin is compact, hypoacetylated, and transcriptionally inactive. How transcription activates upon cell-cycle re-entry is undefined. Here we report robust, widespread transcription within the first minutes of quiescence exit. During quiescence, the chromatin-remodeling enzyme RSC was already bound to the genes induced upon quiescence exit. RSC depletion caused severe quiescence exit defects: a global decrease in RNA polymerase II (Pol II) loading, Pol II accumulation at transcription start sites, initiation from ectopic upstream loci, and aberrant antisense transcription. These phenomena were due to a combination of highly robust Pol II transcription and severe chromatin defects in the promoter regions and gene bodies. Together, these results uncovered multiple mechanisms by which RSC facilitates initiation and maintenance of large-scale, rapid gene expression despite a globally repressive chromatin state.

## Introduction

For decades scientists have used budding yeast to uncover mechanisms of chromatin regulation of gene expression; and the vast majority of these studies were performed in exponentially growing (hereafter log) cultures [1]. Log phase, however, is not a common growth stage in unicellular organism lifecycles. Furthermore, many cell populations in multicellular organisms, such as in humans, are not actively dividing [2–4]. Indeed, the majority of “healthy” cells on Earth are not sustained in a persistently dividing state [3]. Non-proliferating cells reside in a G_0_ state, which generally means these cells are either terminally differentiated, senescent, or quiescent. The quiescent state provides advantages to organisms: quiescence allows cells to remain dormant for long periods of time to survive harsh conditions or to prevent over-proliferation [3–5,2]. Notwithstanding this so-called “dormant state”, quiescent cells can exit quiescence and re-enter the mitotic cell-cycle in response to growth cues or environmental stimuli, which distinguishes quiescence from other G_0_ states. A major hallmark of quiescence is the chromatin landscape—vast histone de-acetylation and chromatin compaction occur during quiescence entry [6–8]. These events happen alongside a global narrowing of nucleosome depleted regions (NDR) and increased resistance to micrococcal nuclease (MNase) digestion, indicating a repressive chromatin environment [6]. Together, these features of quiescent cells point to a critical role for chromatin regulation of the quiescent state. However, the role of chromatin regulation upon exit from quiescence is unknown.

Reversibility is a conserved hallmark of quiescent cells and is required for proper stem-cell niche maintenance, T-cell activation, and wound healing in metazoans [4, 9]. Therefore, we sought to elucidate molecular mechanisms by which cells can overcome this repressive chromatin environment to re-enter the mitotic cell cycle. Given its genetic tractability, the ease by which quiescent cells can be purified, and high level of conservation among chromatin and transcription machinery, we turned to the budding yeast *Saccharomyces cerevisiae* [10]. We can easily isolate quiescent yeast cells after seven days of growth and density-gradient centrifugation. In this context, we can study pure populations of quiescent yeast, a cell fate that is distinct from other cell types present in a saturated culture [11].

Since DNA is wrapped around an octamer of histone proteins in increments of ∼147bp to form nucleosomes [12], enzymes must move nucleosomes to give access to transcription initiation factors [13]. One such enzyme is the SWI/SNF-family member, RSC, which is a 17-subunit chromatin remodeling enzyme complex [14]. RSC contains an ATP-dependent translocase, Sth1 [15–18], multiple subunits with bromodomains (more than half of all bromodomains in the yeast genome are in RSC) and two zinc-finger DNA-binding domains, which allow RSC to target and remodel chromatin [19, 20]. Many components of the RSC complex are essential for viability in budding yeast and the complex is conserved in humans, where it is named PBAF. In humans, mutations in PBAF genes are associated with 40% of kidney cancers [21]; and 20% of all human cancers contain mutations within SWI/SNF family genes [22], underscoring the importance of such complexes in human health.

The best-described role for RSC in regulating chromatin architecture and transcriptions is its ability to generate NDRs, by sliding or evicting nucleosomes [23–25]. Moving the +1 nucleosome allows for TATA binding protein (TBP) promoter binding and transcription initiation [26]. To this end, RSC mostly localizes to the -1, +1, and +2 nucleosomes in log cells [27–29]. However, RSC has also been implicated in the transcription elongation step where it tethers to RNA polymerase and can localize to gene bodies [30–32]. Additionally, RSC binds nucleosomes within the so-called “wide NDRs”, where there are MNase-sensitive nucleosome-sized fragments, known as “fragile” nucleosomes [33–36]. These RSC-bound nucleosomes are likely partially unwrapped to aid in rapid gene induction [36–39].

In this study, we investigated how genes are transcribed during the first minutes of quiescence exit. We were particularly interested in uncovering mechanisms to overcome highly repressive chromatin found in quiescent cells. Unexpectedly, ∼60% of the yeast genome was transcribed by RNA polymerase II (Pol II) by the first 10-minutes of exit, despite the highly repressive chromatin architecture present in quiescence. We found that this hypertranscription [40] event is RSC dependent and that RSC binds across the genome to ∼80% of NDRs in quiescent cells. Upon RSC depletion, we observed canonical abrogation of transcription initiation, defects in Pol II clearance past the +1 nucleosome, and gross Pol II mislocalization, resulting in abnormal upstream initiation and aberrant non-coding antisense transcripts. We further showed that RSC alters chromatin structure to facilitate these processes. Taken together, we propose a model in which RSC is bound to NDRs in quiescent cells to facilitate robust and accurate burst of transcription upon quiescent exit through multiple mechanisms.

## Results

### Hypertranscription occurs within minutes of nutrient repletion post-quiescence

To determine the earliest time at which transcription reactivates during quiescence exit, we fed purified quiescent cells YPD medium and took time points to determine the kinetics of Pol II C-terminal domain (CTD) phosphorylation by western blot analysis (Fig. 1A). Unexpectedly, Pol II CTD phosphorylation occurred within three minutes (Fig. 1A, compare lanes 1 and 2), which was our physical limit of isolating cells during this time course. To determine which transcripts were generated during these early quiescence exit events, we performed nascent RNA-seq using 4-thio-uracil (4tU) to metabolically label new transcripts [41, 42]. In agreement with the western-blot analysis, we observed a high level of transcriptional activation within a few minutes of nutrient repletion (Fig. 1B). Based on our western-blot result, the highest Pol II CTD phosphorylation is observed ∼ten minutes after refeeding. Consistent with this result, we observed the highest level of nascent transcripts at the ten-minute time point, where ∼1000 mRNAs (20% of the genome) were statistically significantly increased compared to the zero-minute time point (Fig.1B, Fig.1—supplement 1A). Given how quickly Pol II was phosphorylated and transcripts were generated, we sought to determine if high levels of Pol II were already bound to the early exit genes in the quiescent state, as was observed previously in a heterogenous population of stationary phase cells [43]. To this end, we performed spike-in-normalized ChIP-seq analysis of Pol II in quiescent cells and at several time points following refeeding (Fig. 1C, Fig. 1—supplement 1B). Low Pol II occupancy levels (compare heatmaps 1 and 5) were detected in quiescent cells, which agrees with our western blot and RNA-seq analyses and previously published literature [6–8]. This implied that Pol II is not paused (Fig. 1C, compare heatmaps 1 and 2) in quiescent cells, and suggested that Pol II needs to be recruited *de novo* for rapid initiation and elongation. In support of this conclusion, we detected only low levels of the pre-initiation complex subunit TFIIB bound to genes in quiescent cells, which increased ∼3-fold by five minutes of exit (Fig. 1—supplement 1C).

**Figure 1.**
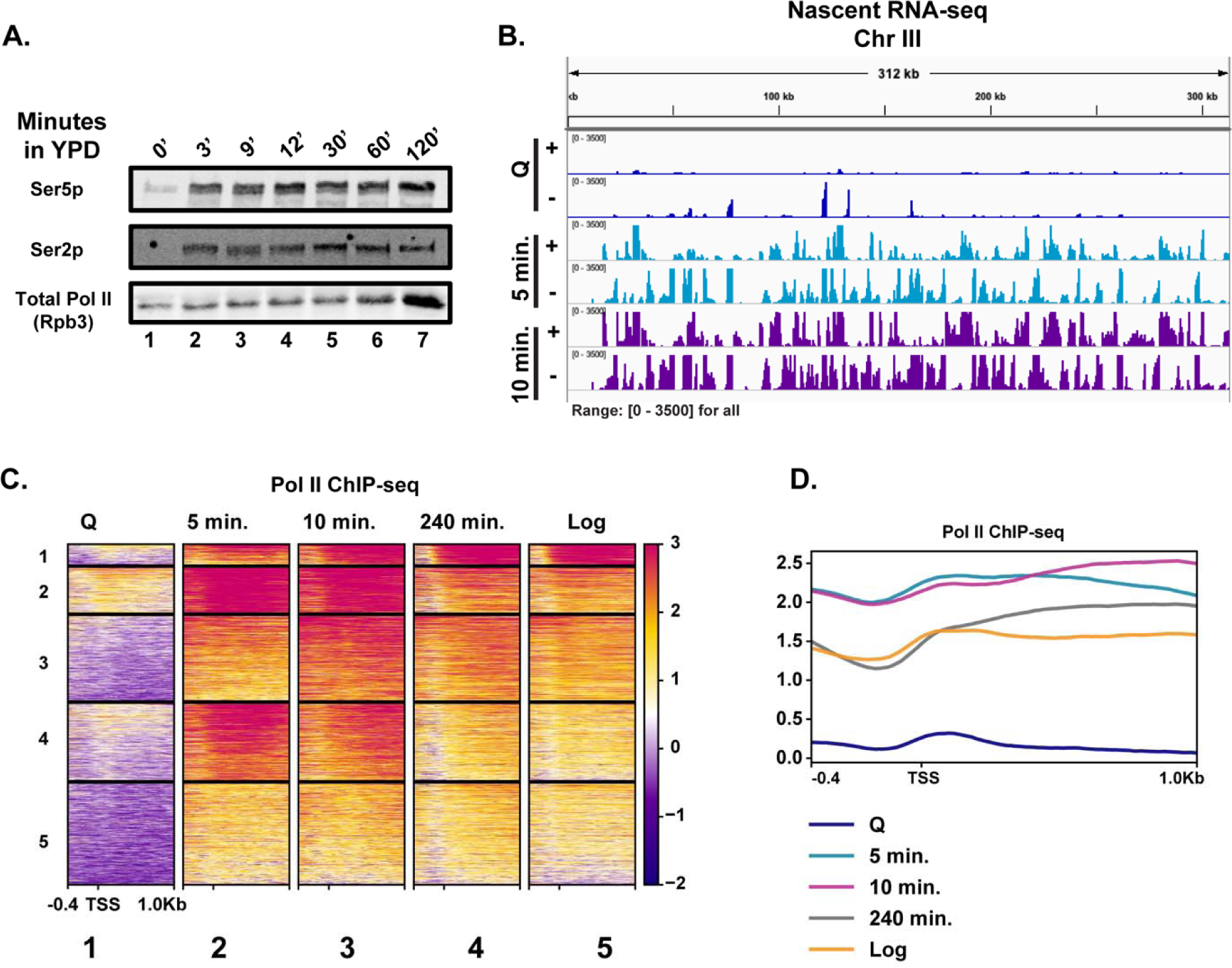
Rapid hypertranscription occurs upon nutrient repletion of quiescent cells. **(A)** Western blots were probed with antibodies to detect Ser5p and Ser2p of the CTD of Rpb1 subunit of Pol II. An antibody against the Rpb3 subunit of Pol II was used as a loading control. **(B)** Strand-specific 4tU-seq analysis. “+” indicates Watson strand and “-“ indicates Click strand. **(C)** Pol II ChIP-seq analysis. Heatmaps show k-means clusters of 6030 genes. Genes are linked across the heatmaps. **(D)** Metaplots of ChIP-seq data shown in (C) without k-means clustering.

Highlighting the high level of transcription occurring in the first ten minutes of quiescence exit, we observed a drop-off in Pol II occupancy levels around the first G2/M phase (240 minutes) (Fig. 1C-D, Fig. 1—supplement 1D). Indeed, when the data were sorted into k-means clusters across the time course, we noticed that many of the genes expressed in the 240-minute time point were similar, but still not identical, to those expressed in log cells, suggesting a recovery to log-like gene expression profile takes hours post refeeding (Fig. 1C, compare columns 4 and 5, Fig. 1D). There was a ∼1.7-fold increase in overall Pol II occupancy in the 10-minute time point relative to that of log cells (Fig. 1D, Fig.1—supplement 1B). Together, these results demonstrate transcription activates extremely rapidly and robustly in response to nutrient repletion.

### Chromatin bears hallmarks of repression during early quiescent exit time points

Given the exceptionally high transcriptional response during the first ten minutes of quiescence exit, we wondered whether chromatin changes reflected hypertranscription. To this end, we performed ChIP-seq analysis of H3 to measure nucleosome occupancy levels genome wide over time. Global H3 patterns during the early exit time points, especially at the 5-minute time point, were more similar to that of the quiescent state than to the 240-minute time point (Fig. 2A, compare columns 1-3), despite higher transcription levels. The most striking changes in histone occupancy during the early time-points were within NDRs, where the pattern at the 10-minute timepoint resembles the 240-minute time point (Fig. 2A, B). However, the H3 profiles outside of NDRs (Fig. 2A, compare column 1-3 and 4 to the right of NDR, and Fig.2B) remain similar to that of quiescent state during the early stage of quiescent exit. In addition to nucleosome occupancy, we tested nucleosome positioning using MNase-seq analysis where nucleosomes with 80% of the digested chromatin is represented by mononucleosomes. Globally, nucleosome positions were stable across the early exit time points (Fig. 2C).

**Figure 2.**
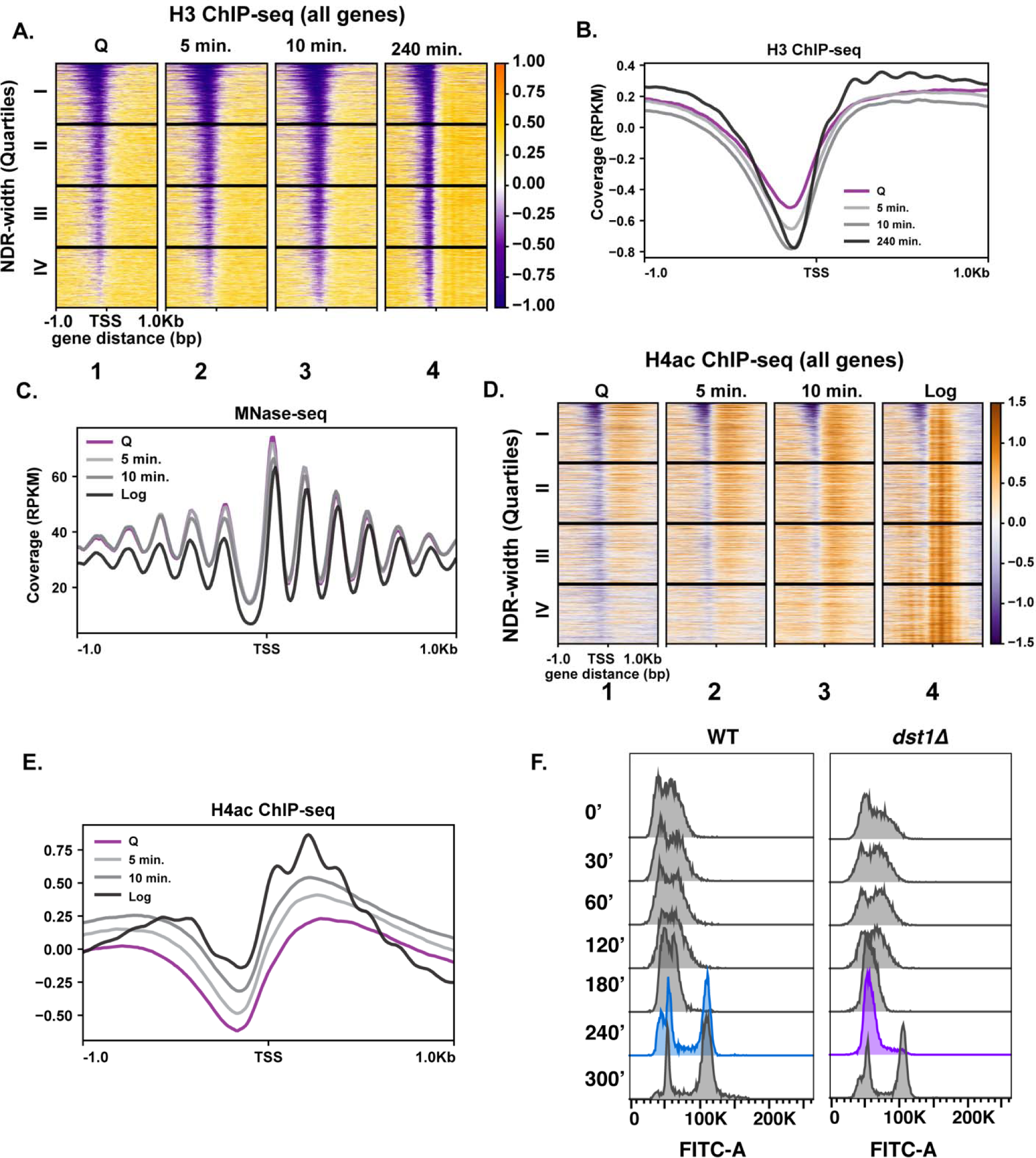
Repressive chromatin persists during early quiescence exit. **(A, B)** ChIP-seq of total H3 in quiescent cells and exit time points sorted into quartiles based on NDR width. **(C)** MNase-seq analysis of 6030 genes in Q (pink line), Log (black line), and Q-exit time points 5 minutes (light grey line) and 10 minutes (dark grey line). **(D, E)** ChIP-seq analysis of penta-acetylated H4 (H4ac) in Q and Log cells and exit time points. Genes are separated as in (B). **(F)** DNA content FACS analysis following Q exit in WT and a TFIIS-absent strain (*dst1*Δ).

We next tested if a burst of histone acetylation occurred during these early exit time points to help overcome the repressive quiescent chromatin environment. To test this, we performed ChIP-seq analysis of H4ac using an antibody that recognizes penta-acetylated H4. Similar to nucleosome occupancy and positions, a modest increase in histone H4 acetylation occurred, but the levels did not reflect that of log cells (Fig. 2D, E). This suggests that, while there was a strong transcriptional response during refeeding, histone acetylation was delayed. This is consistent with a previous study of a mixed population of saturated cultures where histone acetylation was found to occur later in exit[44]. Together, our results are in agreement with a recent study demonstrating that histone acetylation takes place mostly as a consequence of transcription [45].

To assess a biological readout of the repressive chromatin environment, we turned to phenotypic analysis of TFIIS disruption. TFIIS is a general elongation factor that rescues stalled Pol II; and nucleosomal barriers have been shown to increase stalled Pol II [46]. Given that Pol II stalling is common across the genome [47], it is paradoxical that the gene encoding TFIIS is not essential for viability in actively dividing cells, and its deletion does not cause strong growth defects [48]. Since Pol II must achieve a high level of transcription in the repressive chromatin environment during early quiescence exit, we hypothesized that TFIIS may play more critical roles during this period than during log culture. Indeed, in the absence of TFIIS (*dst1*Δ), quiescent yeast cells exhibited defects in cell cycle re-entry, where cells lacking TFIIS stall at the first G1 during exit, which is not the case during the mitotic cell cycle (Fig. 2F, Fig. 2—supplement 1B). These results collectively revealed that the chromatin environment remains repressive during early quiescence exit.

### In quiescence, RSC re-localizes to NDRs of genes expressed in exit

Given the modest changes in chromatin at most genes during the early stage of quiescence exit (Fig. 2), we wondered whether MNase-sensitive or “fragile” nucleosomes were present at the promoters of rapidly induced genes in quiescence and were removed in early exit. Thus, we performed a weaker (low) MNase digestion (10% mononucleosomes) (Fig. 3A) and compared it to the stronger (high) MNase digestion (80% mononucleosomes) (Fig. 3B). Supporting our hypothesis, comparing the weaker MNase digest to the stronger MNase digest revealed that ∼1000 genes have fragile nucleosomes in quiescent cells, which are reduced during exit (Fig. 3A). Additionally, we noticed that the MNase-sensitivity of the +1 and +2 nucleosomes increased at the 10-minute time point, likely coinciding with transcription.

**Figure 3.**
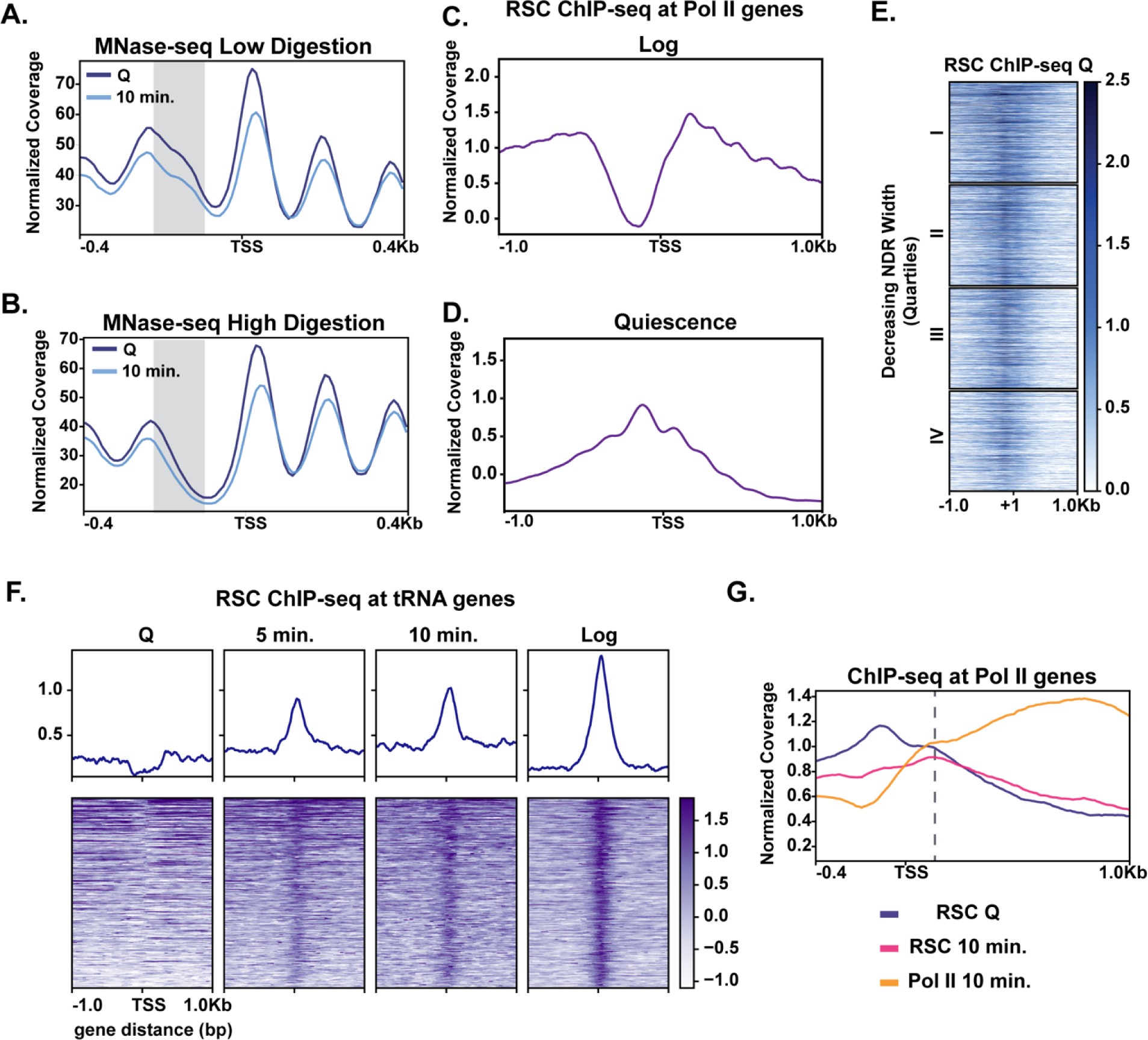
MNase sensitivity and quiescence-specific RSC relocalization indicate remodeling activity required for early exit. **(A)** MNase-digested chromatin to 10% mononucleosomes (low digestion). **(B)** Metaplot of MNase-digested chromatin to 80% mononucleosomes (high digestion) in Q and 10-minute time points. **(C,D)** ChIP-seq of the catalytic RSC subunit in quiescent and log cells at Pol II-transcribed genes. **(E)** ChIP-seq analysis of RSC shown across quartiles based on MNase-seq determined NDR width. **(F)** ChIP-seq of RSC at tRNA genes. **(G)** ChIP-seq of RSC and Pol II comparing RSC movement with Pol II into gene bodies.

It has been recently suggested that that the ATP-dependent chromatin remodeler RSC can remove fragile nucleosomes from promoters to activate transcription [26]. Additionally, it was proposed that RSC-bound nucleosomes are remodeling intermediates that render such nucleosomes more MNase-sensitive [38]. Thus, RSC was a strong candidate for regulating rapid transcription activation during quiescence exit. We performed ChIP-seq analysis of the RSC catalytic subunit Sth1 in quiescent cells. In quiescence, Sth1 exhibited a striking difference in binding pattern compared to log cells (Fig. 3C, D). Sth1 bound to ∼80% of NDRs at gene promoters in quiescent cells (Fig. 3E, Figure 3—supplement 1A). This result was distinct from log cells, where RSC was reported to occupy the widest NDRs but otherwise bind the -1, +1, and +2 nucleosomes for most highly expressed genes (Fig. 3C) [28,26,38]. The RSC binding pattern in quiescent cells instead mirrored a recently described binding pattern in heat shock, where RSC and other transcription regulators *transiently* relocate to the NDRs [49]. In contrast to the heat shock response, however, we observed a stable, strong binding pattern of RSC in NDRs regardless of NDR width (Fig. 3E). Another obvious distinction of RSC binding patterns between log and quiescence was observed at tRNA genes (Fig. 3F). RSC’s role at tRNA expression has been well-studied in log cells [50–52]. In quiescence, RSC was occluded from tRNAs genes. Whereas upon exit, RSC rapidly targeted tRNAs, mimicking the log pattern. Together these data suggest that RSC adopts a quiescence-specific binding profile, one in which RSC is bound to NDRs more broadly across the genome.

We next sought to gain insight into how quiescent RSC occupancy patterns might predict Pol II occupancy during exit. To this end, we compared localization of RSC and Pol II in quiescence and exit. We first found that the presence of RSC at NDRs in quiescent cells and strong transcription in exiting cells co-localized (Fig. 3— supplement 1A). Next, we examined RSC occupancy changes during quiescence exit at Pol II-transcribed genes. During quiescence exit, RSC began to move out of NDRs and into gene bodies as transcription increased (Fig. 3G). These results suggested that RSC facilitates transcriptional activation upon exit and raised the possibility that RSC binding in NDRs may be a mechanism for cells to prepare for quiescence exit.

### RSC depletion causes quiescent exit defects and global Pol II occupancy reduction during quiescence exit

To test the requirement of RSC in quiescence exit, we simultaneously depleted two essential subunits of the RSC complex, Sth1 and Sfh1, using the auxin degron system [53], during quiescence entry (see methods; Figure 3—supplement 1B). Depletion of these subunits throughout the exit process (hereafter “-RSC”) caused a dramatic defect in cell cycle progression upon quiescence exit, where the cells exhibited strong delays in exiting the first G1 stage (Figure 4A). This result contrasted with that in cycling cells, where *rsc* mutants or conditional alleles cause G2/M arrest [54].

**Figure 4.**
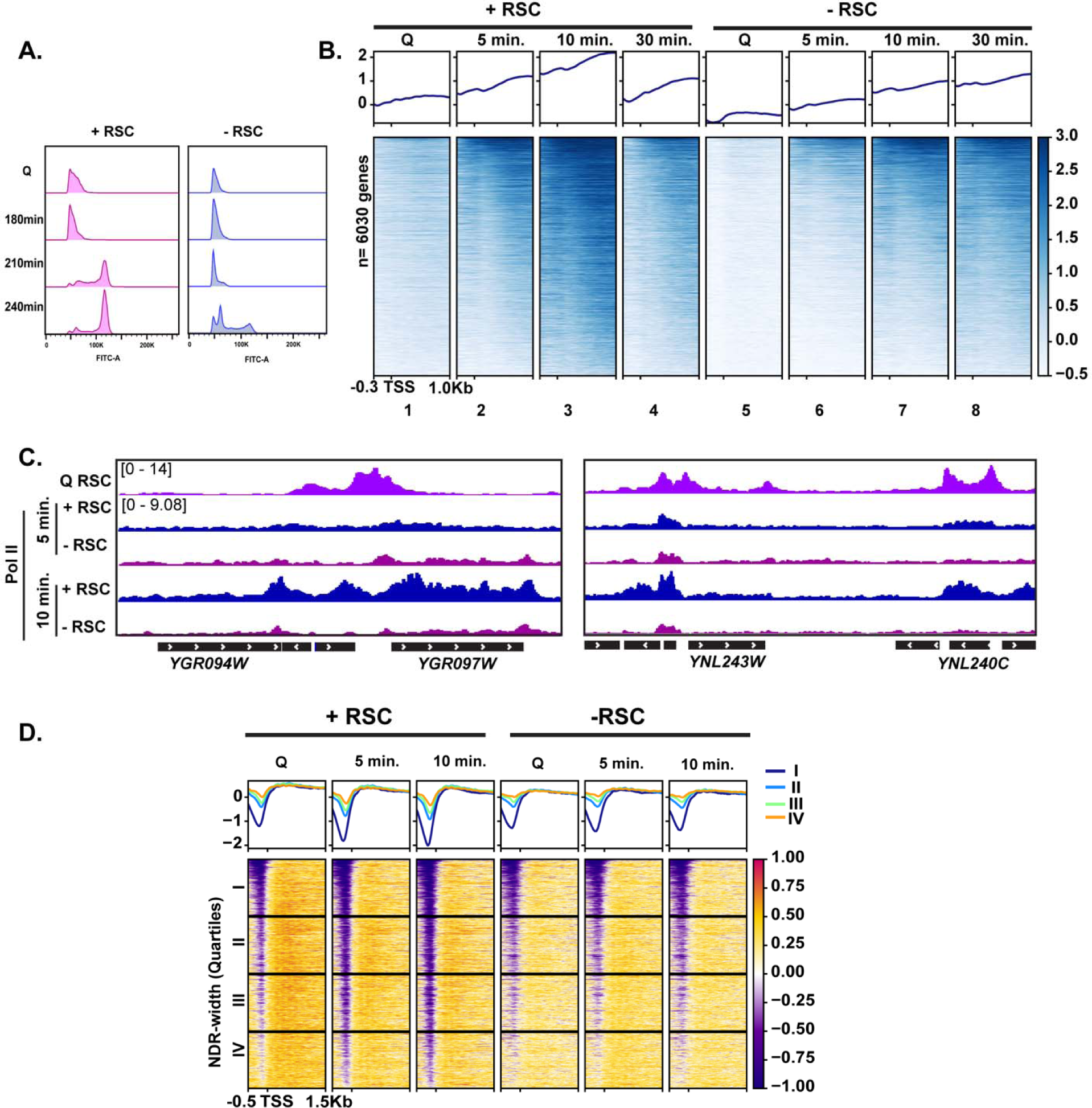
RSC is required for normal quiescence exit and hypertranscription upon nutrient repletion. **(A)** DNA content FACS analysis indicating cell cycle progression during Q exit in the presence (+) or absence (-) of RSC. **(B)** ChIP-seq analysis of Pol II across time in the presence or absence of RSC. Genes are sorted the same in all heatmaps. **(C)** Example tracks of data shown in **(B)** with RSC ChIP-seq in Q cells added. **(D)** H3 ChIP-seq sorted by NDR width (as determined by MNase-seq experiments).

To determine the impact of RSC depletion on hypertranscription during quiescence exit, we performed Pol II ChIP-seq analysis on cells exiting quiescence. In the presence of RSC, Pol II levels peaked at 10 minutes and substantially decreased at 30 minutes after the exit (Fig. 4B, compare columns 3 and 4). As is the case in log cultures [50,55,56], Pol II occupancy decreased in the absence of an intact RSC complex in Q-cells and upon nutrient repletion thereafter (Fig. 4B). Pol II occupancy did eventually increase over time in the RSC-depleted samples. However, even after 30-minutes, Pol II did not reach the peak level of occupancy seen at the 10-minute mark in the +RSC condition (Fig. 4B, compare heatmaps 3 and 8, and 4C). This suggests that the defect in Pol II occupancy during quiescence exit was not solely due to slower kinetics.

As shown earlier in Figure 3G, we observed RSC leaving the NDRs and moving into gene bodies during quiescence exit. Therefore, we examined the impact of RSC depletion on nucleosome occupancy and positioning. H3 ChIP-seq showed that RSC is required for removal of histones within NDRs (Fig. 4D), which is consistent with RSC’s role as the “NDR creator” [24]. Together, these data provide mechanistic explanations for how RSC facilitates Pol II loading during early stages of quiescence exit.

### RSC is required for Pol II passage through gene bodies

Given that RSC moves from NDRs into gene bodies during quiescence exit (Fig. 3G), we next tested whether RSC could aid transcription after initiation. To this end, we selected ∼2000 genes where RSC moved toward gene bodies and examined RSC localization at the 10-minute time point of quiescent exit. This analysis showed uniform movement of RSC from NDR into gene bodies (Fig. 5A). We next tested whether this RSC movement is dependent on Pol II transcription. To this end, we performed Sth1 ChIP-seq analyses during quiescence exit in the presence of a transcription inhibitor 1,10-phenanthroline (Fig. 5B, Pol II control in Fig. 5—supplement 1A). This experiment demonstrated that the movement of RSC from NDRs into gene bodies was strongly inhibited by 1,10-phenanthroline, establishing that RSC re-localization during quiescent exit is dependent on Pol II transcription.

**Figure 5.**
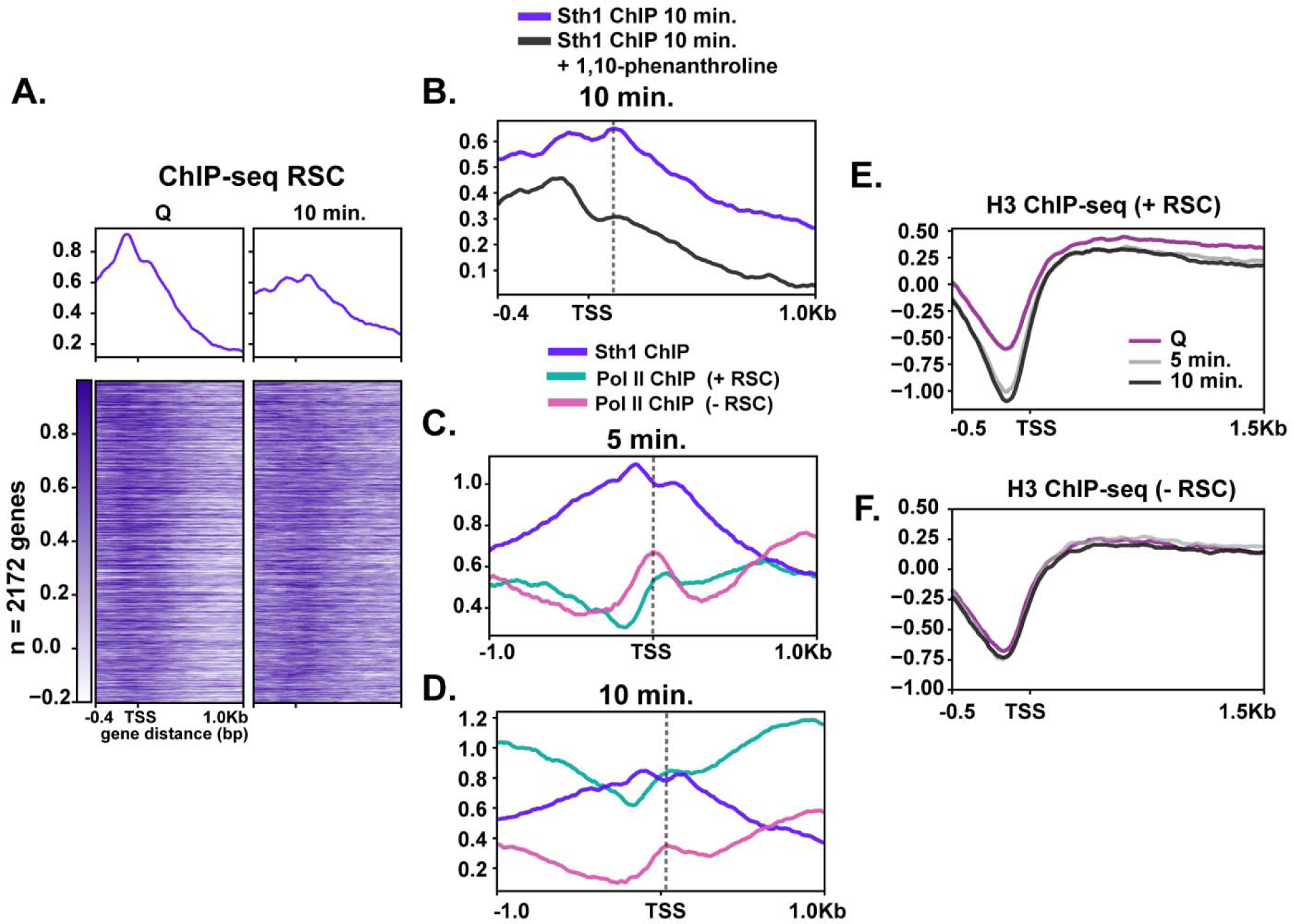
RSC depletion causes severe Pol II mislocalization defects during quiescence exit. **(A)** ChIP-seq of RSC in Q and 10-minute time points. Genes are linked. **(B)**. ChIP-seq of RSC at 10-minutes of exit in the presence and absence of the transcription inhibitor 1,10-phenanthroline. **(C, D)** ChIP-seq of RSC and Pol II during exit. **(E-F)** H3 ChIP-seq in quiescence and during exit in the presence and absence of RSC.

Co-transcriptional movement of RSC into gene bodies suggested a possibility that RSC may help Pol II passage through gene bodies. To test this, we determined the effects of RSC depletion on Pol II localization during early time points of quiescent exit. Fig. 5C and D show that RSC depletion affects Pol II localization in at least two ways during early quiescent exit. First, consistent with Fig 4B, the robust increase in the amount of Pol II over genes is strongly decreased upon RSC depletion. In addition, upon RSC depletion, Pol II sharply accumulates at TSSs at the 5-minute mark, which continued to the 10-minute mark. In sharp contrast, PoI II accumulates at slightly more downstream at the 5-minute mark and moves mostly to downstream regions at the 10-minute time point in the presence of RSC. We also noticed a pile-up of Pol II at the 3’-end of genes at the 5-minute timepoint upon RSC depletion (Fig. 5C). This is in agreement with the possibility that RSC may be involved in proper transcription termination [57]. At these loci, NDRs are relatively shallow in quiescence but histone density rapidly decreases upon quiescence exit in the presence of RSC (Fig. 5E, Fig. 5—supplement 1B). In the absence of RSC at these sites, however, histone density is unexpectedly lower at NDR in quiescence but does not change during quiescent exit (Fig. 5F, Fig. 5—supplement 1B), suggesting defective chromatin structure at and downstream of the NDR. Together, these results are consistent with the notion that co-transcriptional movement of RSC facilitates passage of Pol II through nucleosomes immediately downstream of TSSs through chromatin regulation.

### RSC suppresses abnormal upstream transcription initiation

The fact that Pol II accumulated upstream of TSSs at the 5-minute mark upon RSC depletion (Fig. 5C) suggested possible defects in transcription start site selection. To test this possibility, we examined the 4tU-seq profiles of a subset of RSC targets (1426 genes) in which there appeared to be an enrichment of RNA signal directly upstream and downstream of TSSs. We took the log_2_ ratio of RNA signal in the depleted condition versus the non-depleted condition at the ten-minute time point (Fig. 6A). We sorted the genes using k-means clusters and took into account RSC binding when determining gene sets to analyze.

**Figure 6.**
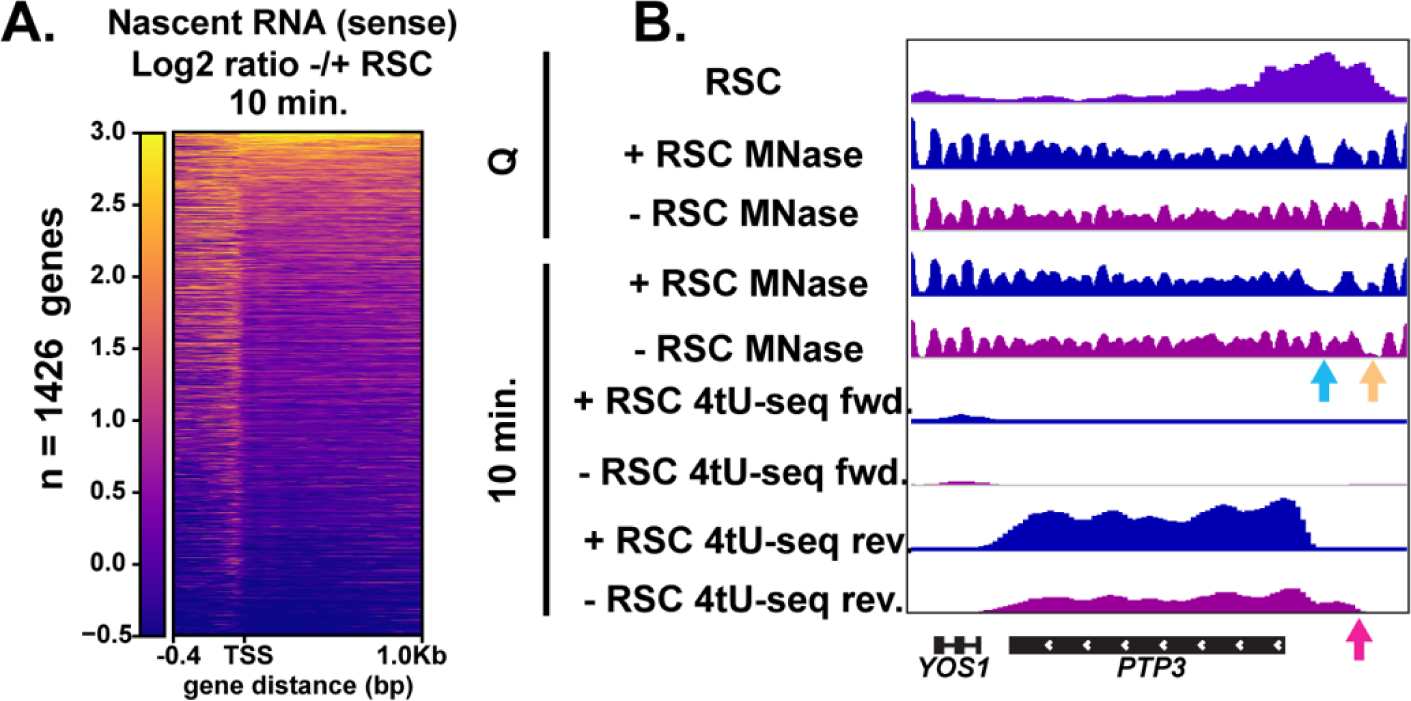
RSC depletion causes upstream transcription relative to canonical TSS. **(A)** Heatmap showing the Log_2_ ratio of nascent sense transcripts in RSC-depleted versus non-depleted cells. Shown are 1426 genes that have upregulated transcripts at the 5’-ends of genes in the sense direction and have RSC ChIP signals. **(B)** Example gene of aberrant upstream transcript. Arrows direct to defects: blue arrow points to loss of NDR, yellow arrow points to gain of NDR, and pink arrow points to upstream RNA signal.

This analysis revealed that upon RSC depletion a large number of genes (∼1400) exhibited increased nascent sense-strand RNA signals starting upstream of their normal TSSs, demonstrating wide-spread defects in TSS selection. Examination of individual loci revealed that, in addition to filling of an NDR at the normal TSSs, an NDR is created upstream, which overlaps with ectopic transcription observed at an upstream TSS (see Fig. 6B for an example). These results suggest that RSC facilitates selection of accurate transcription initiation sites through proper NDR formation upstream of protein coding genes during the burst of transcription during quiescence exit. This is likely a quiescence-specific function of RSC, or a result of the robust hypertranscription event during exit, as depletion of Sth1 in cycling cells mostly repressed transcription initiation with relatively few new upstream transcription start sites [55, 56].

### RSC is required for suppression of anti-sense transcripts during quiescence exit

Given the robust transcriptional response during the early minutes of quiescence exit (Fig. 1), we examined whether aberrant transcripts might also arise during quiescence exit when RSC was depleted. Indeed, in many cases we found antisense transcripts arising in the absence of RSC. We found ∼900 RSC targets that had generally reduced sense transcript levels with strongly upregulated cognate antisense transcripts (Class I), and ∼600 genes (Class II) with only modest changes in both sense and anti-sense transcript levels upon RSC depletion (Fig. 7A). Chromatin analyses of individual Class I loci revealed that RSC depletion caused narrower NDRs upstream of the sense TSS (see Fig. 7B for an example). In contrast, NDRs for sense transcripts remained largely open at Class II genes (Fig. 7B). Both classes have RSC bound at the promoters of the sense genes in quiescence, with slightly higher RSC binding in the class I genes (Fig. 7C). Strikingly, nucleosome positioning was heavily impacted in the Class I set of genes upon RSC depletion in the sense direction, where NDRs became more resistant to MNase and nucleosomes in gene bodies were shifted toward the 5’-ends of genes. This was in contrast to that of Class II where NDRs were largely open (Fig. 7D). Consistent with the MNase-mapping data, nucleosome occupancy at NDRs and in gene bodies are much more strongly affected by RSC depletion at Class I genes than Class II genes (Fig. 7E). It is likely that Class II genes overcome the absence of RSC by having more “fragile” nucleosomes that can be readily removed by general regulatory factors [26]. These results collectively showed that chromatin structure at the Class I genes is especially dependent on RSC. In both classes of genes, RSC signals and RSC-dependent chromatin changes are not apparent around the start sites of anti-sense transcripts. Therefore, suppression of anti-sense transcripts is unlikely to be a direct role for RSC. Instead, it is likely that both Class I and II genes, especially the former, have an intrinsic property to allow anti-sense transcription to occur when not properly regulated, and RSC is targeted to them to ensure sense transcription takes place through formation of proper NDRs.

**Figure 7.**
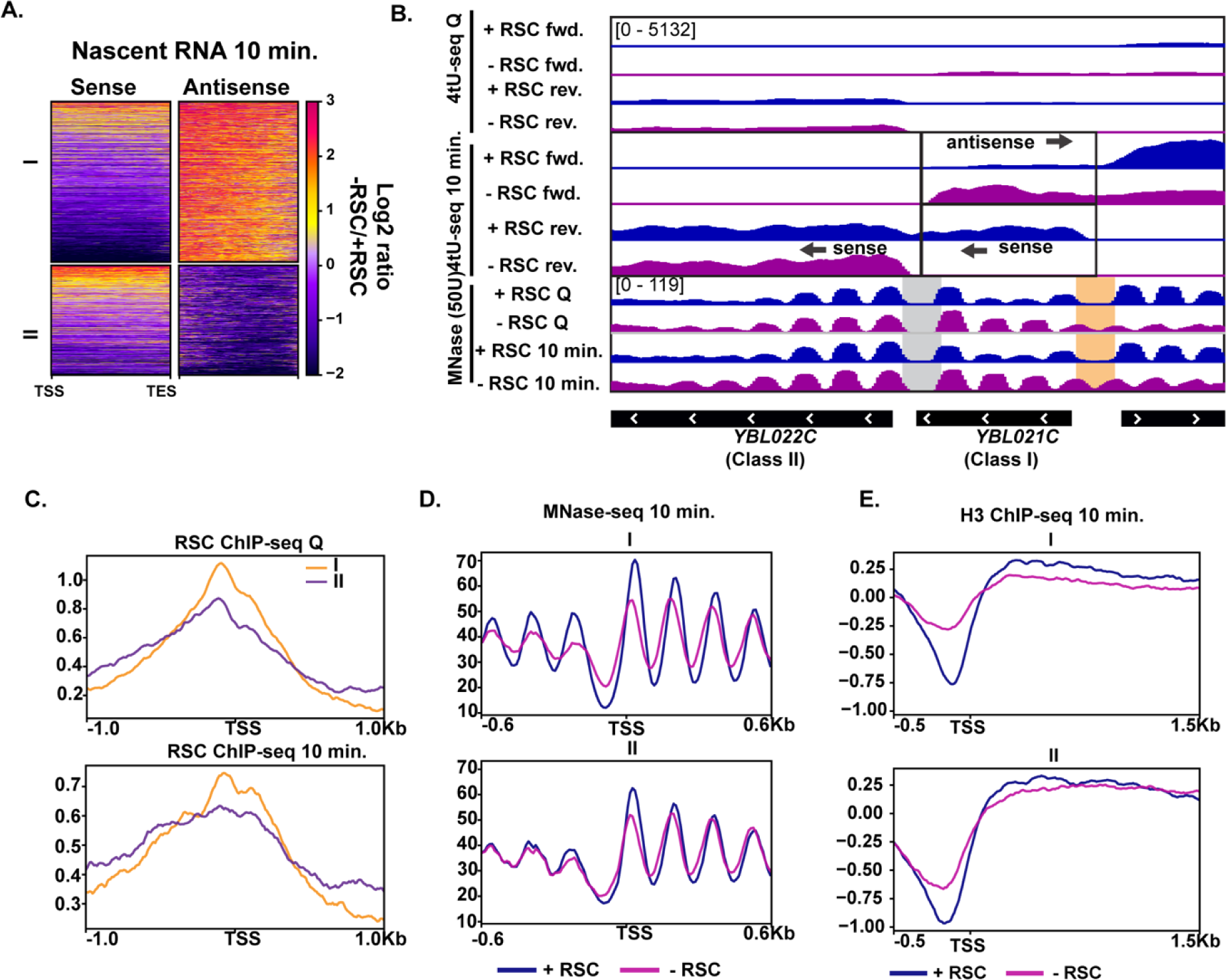
Aberrant antisense transcription arises when chromatin around sense transcripts is abrogated in the absence of RSC. **(A)** Heatmaps of the Log_2_ ratio of nascent RNAs that are RSC targets and give rise to antisense transcripts (cluster I, ∼890 genes) when RSC is depleted and those where antisense transcripts are not made when the sense transcript NDR is unchanged (cluster II, ∼600 genes). **(B)** Browser tracks of 4tU-seq and MNase-seq data. Boxes highlight defects (orange box) in NDRs and where antisense transcription arises from co-opted, intact NDRs (grey box). **(C)** ChIP-seq of RSC in quiescent cells and during exit at both cluster I and II. **(D)** MNase-seq at the 10-minute time point. (**E**.) 10-minute time point during exit of H3 ChIP-seq separated into the two clusters, I and II.

## Discussion

In this report we have shown that there is a rapid and robust transcriptional response during the very early minutes of quiescence exit (Fig. 8A). This response is greatly dependent on the chromatin remodeling enzyme RSC. We found that RSC promotes transcription at the right place and time in four different ways: 1) RSC promotes transcription initiation by creating NDRs in quiescence and maintaining them during exit (Fig. 8B). 2) RSC moves into gene bodies and helps Pol II transcribe past the +1 nucleosome (Fig. 8C). 3) RSC maintains proper NDR locations to allow for accurate transcription start site selection (Fig. 8D). 4) RSC suppresses cryptic antisense transcription via generating NDRs at the cognate sense genes (Fig. 8E). Together, our results suggest that the massive transcriptional response requires highly accurate nucleosome positioning to allow for cells to exit from the quiescent state.

**Figure 8.**
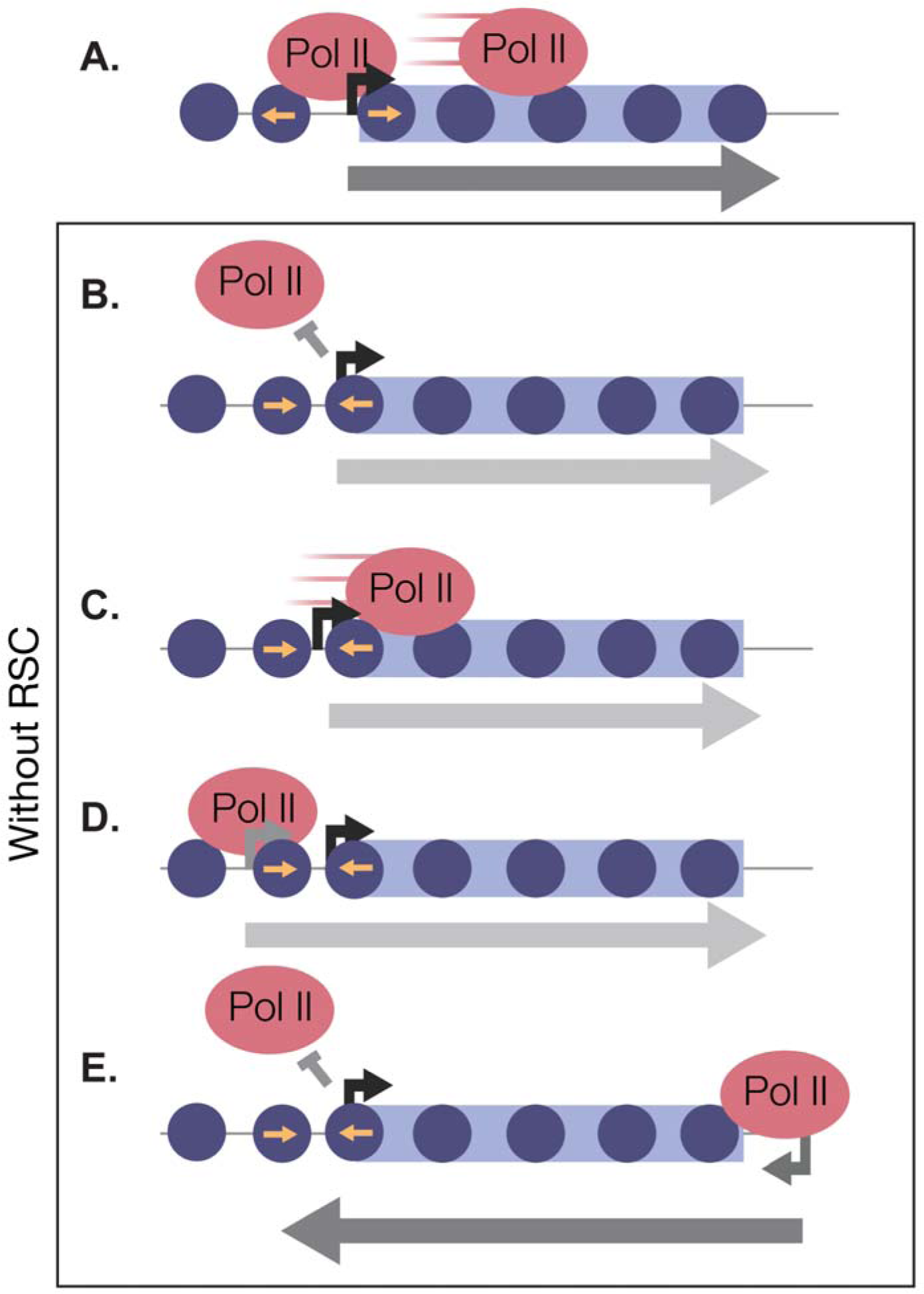
RSC safeguards the quiescent genome from aberrant transcription. In quiescent cells, RSC binds to NDRs upstream of Pol II transcribed genes. Upon quiescence exit, RSC shifts the +1 nucleosome to allow for Pol II occupancy and traverses into gene bodies **(A)**. In the absence of RSC NDRs are globally narrower and transcription initiation is blocked **(B).** At a subset of genes, RSC is required for efficient Pol II passage past the +1 nucleosome **(C)** and prevent upstream TSS selection **(D).** NDRs that are open despite RSC depletion become cryptic promoters and are utilized by transcription machinery to generate aberrant lncRNAs and antisense transcripts **(E)**.

Quiescent yeast must downregulate their transcriptional program and generate a repressive chromatin environment in order to survive harsh conditions for extended time periods [10,6,58,59]. How, then, do cells rapidly escape the quiescent state when conditions are favorable? In this study, we show that there is an incredibly strong transcriptional response to nutrient repletion after quiescence, notwithstanding a relatively repressive chromatin environment that persists until the first G2/M phase after quiescence. Indeed, we identified a previously unidentified phenotype for the deletion of the gene encoding yeast TFIIS, *dst1*Δ. High numbers of stalled Pol II are present in cycling cells [47] despite the little impact of deleting *DST1* on cycling cell growth. We speculate cells exiting quiescence may rely more heavily on TFIIS to transcribe through repressive chromatin [60, 61].

During quiescence, RSC relocates to NDRs upstream of Pol II transcribed genes that are transcribed in exit. Although RSC binds and regulates chromatin around Pol III genes [27, 50], RSC is depleted at tRNA genes in quiescence and only returns during quiescence exit, further supporting the notion that RSC is globally re-targeted in quiescence. This is distinct from the transient NDR-relocalization observed in heat shock [49], as what we observed in quiescence was a sustained and rather stable localization. How RSC binds to these new locations in quiescence is unknown. Given the distinct structure of quiescent chromatin there are several, non-mutually exclusive, explanations for RSC’s binding pattern in quiescence. 1) The genome is hypoacetylated and thus RSC can no longer bind to acetylated nucleosomes in quiescence via its bromodomains [19]. However, given the highly robust response to refeeding, RSC activity must be poised to be active in this state. An intriguing possibility could be that histone acetylation inhibits RSC activity to some extent as was recently reported *in vitro* [62]. This would be consistent with the rapid changes in nucleosome positioning at many genes during quiescence exit in the absence of high levels of histone acetylation. 2) Recent structural studies have shown that the nucleosome acidic patch is in direct contact with subunits of the RSC complex [63–66]. If the acidic patch is occluded by hypoacetylated H4 tails in quiescence for example [12,67–70], it is possible that RSC can no longer interact with this region of the nucleosome, rendering its binding abilities different in quiescence. Finally, 3) a lack of Pol II activity in quiescent cells could prevent RSC from moving out of NDRs and into gene bodies. Indeed, transcription appears to play a prominent role in RSC localization: RSC moves into gene bodies during transcription activation and this movement is blocked when transcription is inhibited, as we have reported above. It is likely that a combination of transcription and histone acetylation helps pull RSC into gene bodies, given recent work showing that acetylation is a consequence of transcription [45]. In a separate study, we recently found that the SWI/SNF remodeling enzyme promotes transcription of a subset of hypoacetylated genes during quiescence entry, implying a specialized transcription regulation program for essential genes in the wake of widespread transcriptional shutdown [58]. In cycling cells, it was recently shown that RSC and SWI/SNF cooperate at a subset of genes [71]. Our results suggested that cooperation between the two SWI/SNF class remodeling factors may also occur during quiescence entry.

Consistent with co-transcriptional re-localization, our data suggest RSC plays an active role in helping Pol II transcribe past the +1 nucleosome in addition to initiating transcription. Supporting this idea was our observation of a subset of genes where RSC depletion caused a Pol II enrichment around the +1 nucleosome. Previous reports showed that RSC can bind gene bodies and impact elongating and terminating Pol II [31, 57]; and one study showed interactions between the Rsc4 subunit and all three RNA polymerases [30]. An intriguing possibility could be that RSC directly interacts with Pol II to facilitate transcription past the first few nucleosomes.

The transcriptional response during quiescent exit was dampened by depleting the essential chromatin remodeler, RSC, but it did not diminish completely. Pol II occupancy was globally decreased ∼2-fold at the 10-minute time point in RSC-depleted cells. However, at ∼900 genes we found that while sense transcription was reduced, antisense transcripts were generated. This was largely due to a nearby NDR susceptible to transcription initiation that could be co-opted for antisense transcription. The mechanism that allows for this cryptic transcription is still unknown. Chromatin remodeling enzymes are vastly important for repressing antisense lncRNAs [72].

Different chromatin remodeling enzymes function to repress lncRNA transcripts in cycling cells, including RSC [73–75]. We speculate RSC is particularly suitable to regulate global transcriptome during quiescence exit due to its high abundance, which allows it to function through multiple mechanisms. The mouse embryonic stem cell-specific BAF complex was also recently shown to globally repress lncRNA expression [76]. This raises the possibility that some of our observations in yeast quiescent cells could be conserved in mammalian quiescent cells. Given the robust transcriptional response that occurs during quiescence exit, it is likely that chromatin structure is crucial for maintaining the quality of the transcriptome. Indeed, we noted cases where transcription occurred upstream of the canonical TSS when an NDR was not generated, highlighting the defects in Pol II initiation and start site selection due to chromatin defects in the absence of RSC. Hypertranscription events similar to the one observed during quiescence exit occur throughout all organisms, particularly during development [40]. Therefore, it is quite possible that we will see similar, multifaceted roles for RSC homologues or other abundant chromatin remodeling factors in facilitating proper hypertranscription in many other systems.

## Materials and Methods

### Yeast strains, yeast growth media, quiescent cell purification, and exit time courses

The *S. cerevisiae* strains used in this study are listed in Supplementary Table S1 and are isogenic to the strain W303-1a with a correction for the mutant *rad5* allele in the original W303-1a [77]. Yeast transformations were performed as previously described [78]. All cells were grown in YPD medium (2% Bacto Peptone, 1% yeast extract, 2% glucose). We note that quiescent (Q) yeast need to be grown in YPD using “fresh” (within ∼three months) yeast extract as a source. To purify Q cells, liquid YPD cultures were inoculated with a single colony into liquid cultures (colonies were no older than one week). Yeast cells were grown in Erlenmeyer flasks ten times the liquid volume for seven days at 30°C and shaking at 180 RPM. Q cells were purified by percoll gradient centrifugation as previously described [11]. Briefly, percoll was diluted 9:1 with 1.5 M NaCl into 25-mL Kimble tubes and centrifuged at 10,000 RPM for 15-minutes at 4°C. Seven-day cultures were pelleted, washed with ddH2O, resuspended in 1 mL of ddH2O, and gently pipetted over a pre-mixed percoll gradient. 400 OD_660_ were pipetted onto a 25-mL gradient. Gradients with loaded cells were centrifuged for one hour at 1000 RPM, 4°C. The upper, non-quiescent cell population and the middle, ∼8 mL fraction, were carefully discarded via pipetting. The remaining volume was washed twice with ddH2O in a 50 mL conical tube at 3,000 RPM, 10 minutes each.

Q exit experiments were performed as follows: Q cells were harvested and added to YPD to 1 OD_660_/mL. Cells were grown at 25°C to slow the kinetics for feasibility. For ChIP-seq and MNase-seq experiments, cells were grown to the appropriate time and then crosslinked for 20 minutes (described in more detail in the sections below).

### Depletion of RSC subunits, Sth1 and Sfh1

The yeast strains YTT 7222 and 7224 were grown in 5-mL overnight YPD cultures, back diluted for four doublings, and inoculated to 0.002 OD_660_ into the appropriate YPD volume for a given experiment. Cells were grown for 16 hours and monitored for glucose exhaustion using glucose strips. Six hours after glucose exhaustion, 1mg/mL of Indole-3-acetic acid (Sigma, I3750-5G-A) was added, in powder form, to the culture. Q cells were purified as described above and depletion efficiency was determined by western blot analysis (Supplementary Figure 1A).

### Western Blot Analysis

Yeast cells were lysed by bead beating in trichloroacetic acid (TCA), as previously described [79]. Proteins were resolved on 8% polyacrylamide gels and transferred to nitrocellulose membranes. Membranes were incubated with primary antibodies: anti-Rpb3 (Biolegend, 665003 1:1000 dilution), anti-Ser5p (Active Motif, 61085 1:1000 dilution), anti-Ser2p (Active Motif, 61083, 1:1000 dilution), and anti-HSV (Sigma, 1:500). Following primary incubation, membranes were incubated with either anti-mouse or anti-rabbit secondary antibodies (Licor, 1:10000). Protein signals were visualized by the Odyssey CLx scanner.

### ChIP-seq

100 OD_660_ U of cells were crosslinked and sonicated in biological duplicate using the protocol described in [80]. Proteins were immunoprecipated from 1 μg chromatin and 1 μL of anti-H3 (Abcam, 1791) conjugated to 20 μl protein G magnetic beads (Invitrogen, 10004D) per reaction. For Pol II ChIPs, we used an antibody against the Rpb3 subunit (2 μl per reaction, Biolegend 665004) conjugated to 20 μl protein G magnetic beads (Invitrogen, 10004D). For Sth1 ChIP experiments we used an antibody against the Flag-epitope tag, FLAG M2 mouse monoclonal (Sigma Aldrich, F1804) and conjugated to 20 μl protein G beads (Invitrogen, 10004D) Libraries were generated using the Ovation Ultralow v2 kit (NuGEN/Tecan, 0344) and subjected to 50-bp single-end sequencing on an Illumina HiSeq 2500 at the Fred Hutchinson Cancer Research Center genomics facility. We used bowtie2 to align raw reads to the sacCer3 reference genome [81]. Reads were then filtered using SAMtools [82]. Bigwig files of input-normalized ChIP-seq data were generated from the filtered bam files using deepTools2 [83] and dividing the IP data by the input data. Matrices for metaplots were generated in deepTools2 using the annotation file from [84].

### MNase-seq

Cell growth and crosslinking was done in the same fashion as in ChIP-seq experiments. Generally, we followed the protocol in [80], with changes described here. Cells were spheroplasted using 10 mg zymolyase (100T, AMSBIO, 120493-1) per 100 OD_660_ cells. For Q cells, zymolyase treatment could take up to two hours. We monitored the cells via microscopy and stopped the spheroplasting step when ∼80% of the cells were spheroplasted. MNase digestion was performed as described in [80]. High digests (80% mononucleosomes) required 50U of micrococcal nuclease (Worthington, LS004798) and for the low digests, chromatin was treated with 10 U of MNase. From this step, chromatin was reverse crosslinked as described in [80]. Following reverse crosslinking, RNase, and proteinase-K digestion, DNA was phenochloroform-extracted. Any large, uncut genomic DNA species was separated out using Ampure beads (Beckman).

Sequencing libraries were generated from the purified DNA using the Ovation Ultralow v2 kit (NuGEN, 0344). Libraries were subjected to 50-bp paired-end sequencing on an Illumina HiSeq 2500 at the Fred Hutchinson Cancer Research Center genomics facility. We used bowtie2 to align raw reads to the sacCer3 genome and filtered reads using SAMtools as described above for ChIP-seq analysis. Bigwig files of input-normalized ChIP-seq data were similarly generated from the filtered bam files using deepTools2 and the MNase option to center the reads around nucleosome dyads. Data represented in the paper were filtered to mononucleosome sizes using deepTools2.

### Nascent RNA-seq

Generally, nascent RNA-seq experiments were performed as described in [85, 42]. For the 0-minute and 5-minute samples, we added 100 and 50 OD_660_ of Q cells, respectively, to YPD containing 5 mM 4-thiouracil (Sigma, 440736-1G). Cells were incubated with 4tU for 5 minutes before pelleting (one minute, 3500 RPM) and flash frozen in liquid nitrogen. For the 10-minute time points, 50 OD units of quiescent cells were released into YPD for 5 minutes before an additional 5-minute incubation with 4tU at a final concentration of 5 mM. All time points were labeled with 4tU for a total of 5 minutes before pelleting and freezing. Total RNA was isolated using Ambion’s RiboPure Yeast Kit (Thermo, AM1926). *S. cerevisiae cells* were lysed in the presence of *Kluvomyces lactis* (*K. lactis*) cells in a 100:1 mixture. RNA was treated with DNAseI according to the TURBO DNase kit (Thermo, AM2238). 40 ug RNA was then biotinylated with MTSEA biotin-XX (diluted in 20% DMF) at a final concentration of 16.4 uM in 20mM HEPES pH 7.4 and 1 mM EDTA at room temperature for 30 minutes. Unreacted MTS-biotin was removed from samples by PCI extraction and resuspended in 100 uL nuclease-free water. Strepavidin beads (Invitrogen 65001) were washed with high-salt wash buffer (100 mM Tris, 10 mM EDTA, 1 M NaCl, 0.05% Tween-20) and blocked for one hour in high-salt wash buffer containing 40 ng/uL glycogen. 40 uL of streptavidin beads were added to the RNA samples and incubated for 15 minutes at room temperature. Beads were washed three times in 1 mL high salt wash buffer and eluted for 15 minutes at room temperature in 50 uL streptavidin elution buffer (100 mM DTT, 20 mM HEPES, 2.7, 1 mM EDTA, 100 mM NaCl, 0.05% Tween-20). The resulting RNA was then purified and concentrated using the Qiagen miRNeasy kit (#217084). Libraries were prepared from 5 ng of RNA using the Ovation SoLo kit (NuGEN/Tecan, custom AnyDeplete; contact Tecan for ordering this kit for yeast). Libraries were subjected to 50-bp paired-end sequencing on an Illumina HiSeq 2500 at the Fred Hutchinson Cancer Research Center genomics facility. We used bowtie2 to align raw reads to the sacCer3 and *K. lactis* (Ensembl ASM251v1) genomes and filtered reads using SAMtools as described above for ChIP-seq analysis. Differential expression analysis was performed using DESeq2 [86]

## Data Availability

All sequencing data are uploading on the NCBI Gene Expression Omnibus under the accession number GSE166789.

## Acknowledgements

We are grateful to members of the Tsukiyama lab and Pravrutha Raman for helpful comments and critical reading of this manuscript. We thank Sarah Hainer and Felix Mueller-Planitz for advice on MNase-seq experiments. We thank Benjamin Martin, Rafal Donczew and Sandipan Brahma for advice and feedback. We thank Mitchell Ellison and Alex Francette for advice about analyzing nascent RNA-seq data. TT was supported by the National Institutes of Health (R01 GM111428 and R35GM139429). CEC was supported by the National Cancer Institute (T32CA009657) and National Institutes of Health (F32GM131554).

---

**Figure 1—supplement 1.**
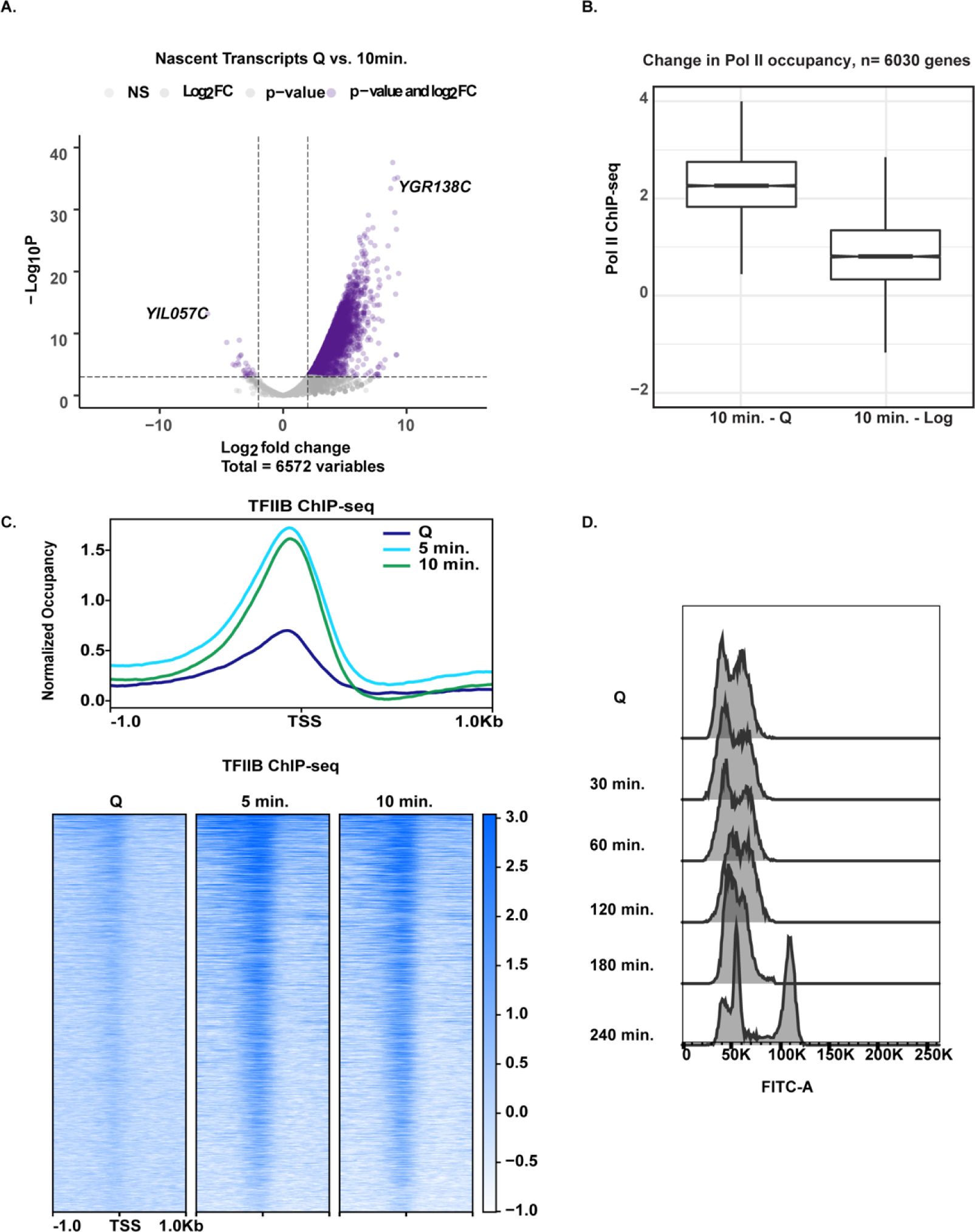
**(A)** Volcano plot of nascent transcripts comparing significant changes in expression using a 2-fold cut off. **(B)** Boxplots illustrating the difference in Pol II ChIP-seq signals across genes. Log_2_ ratio values were subtracted (ex: Q log_2_ values were subtracted from 10 min. log_2_ values). **(C)** TFIIB ChIP-seq analysis in Q cells and exit time points. Genes are linked across the time points and are aligned to TSS. **(D)** DNA content FACS analysis indicating cell cycle progress during Q exit.

**Figure 3—supplement 1.**
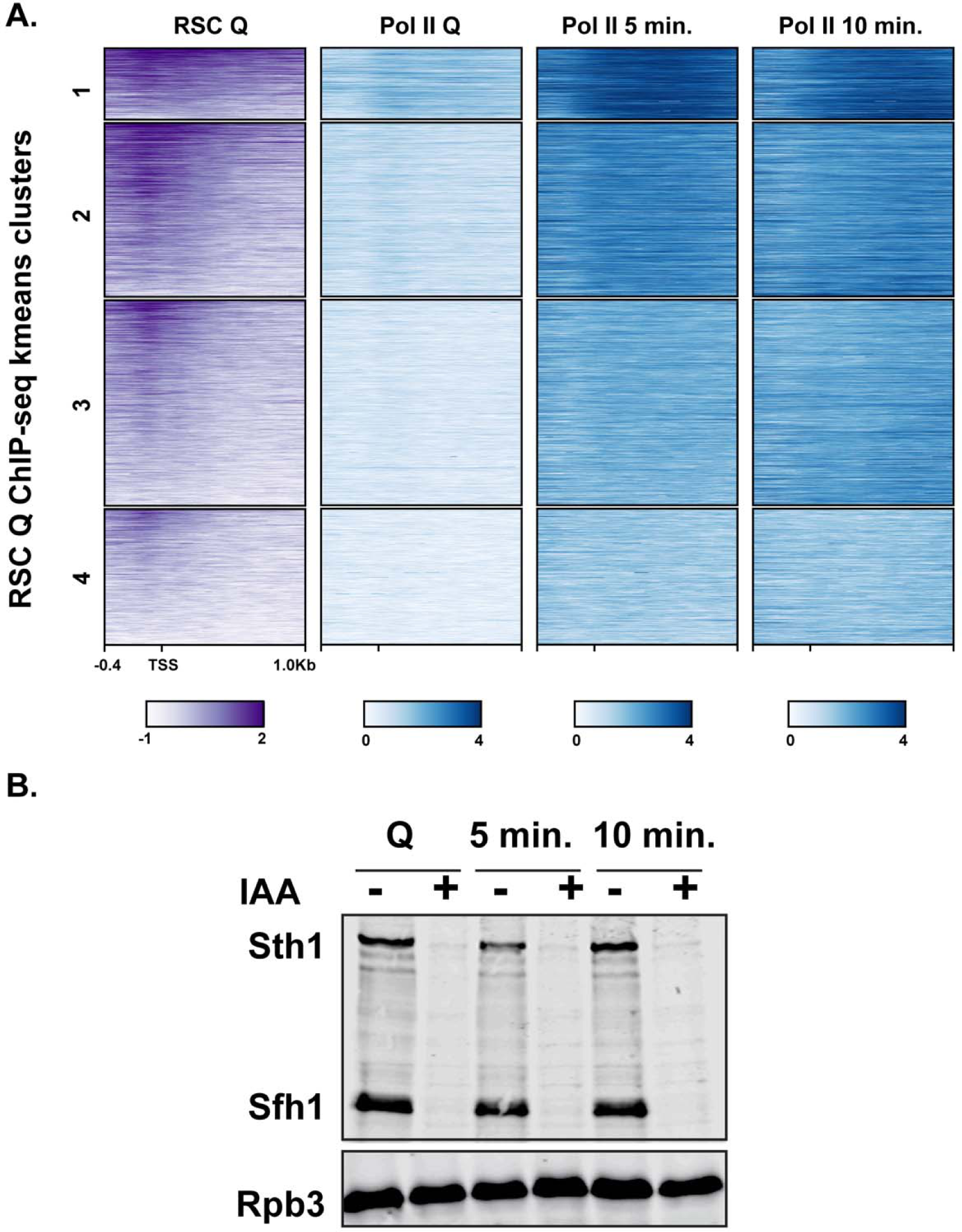
**(A)** ChIP-seq analysis of RSC and Pol II using antibodies against Flag-tagged Sth1 and Rpb3, respectively. Genes are sorted into k-means clustered and are linked across the different ChIPs. **(B)** Western blot analysis of RSC depletion. Both Sth1 and Sfh1 contain C-terminal HSV and AID tags for detection and depletion using IAA. Western blot was probed with an antibody recognizing the HSV epitope tag and Rpb3 (Pol II subunit) as a loading control. The addition of IAA is indicated by – or +.

**Figure 5—supplement 1.**
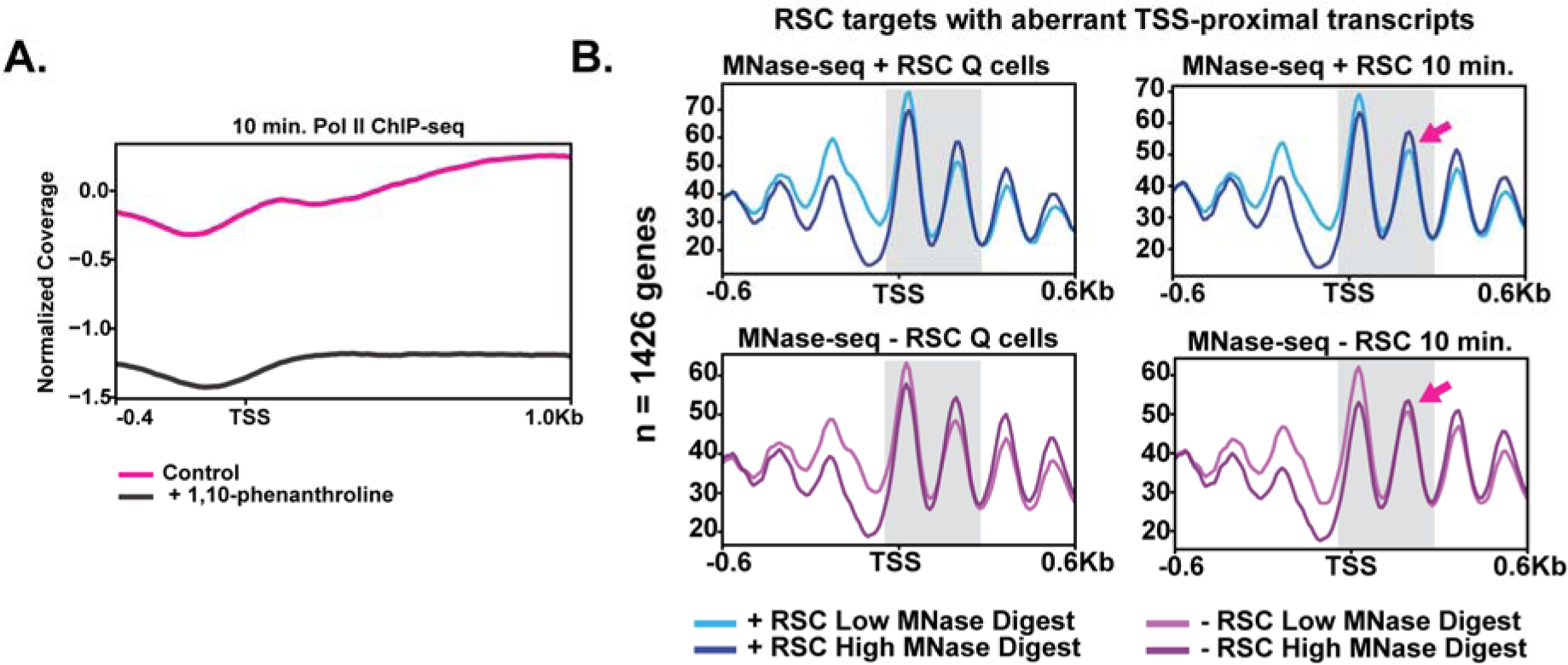
**(A)** ChIP-seq analysis of Pol II in the absence and presence of the transcription inhibitor 1,10-phenanthroline. **(B)** MNase-seq analysis assessing differences in MNase sensitivity in Q and ten-minutes for cells with and without RSC. The +2 nucleosome MNase-digestion differences are highlighted by the pink arrows.

## Notes

### Competing Interest Statement

The authors have declared no competing interest.

